# Evolution of DNA methylation across Ecdysozoa

**DOI:** 10.1101/2021.07.15.452454

**Authors:** Jan Engelhardt, Oliver Scheer, Peter F. Stadler, Sonja J. Prohaska

## Abstract

DNA methylation is a crucial, abundant mechanism of gene regulation in vertebrates. It is less prevalent in many other metazoan organisms and completely absent in some key model species, such as *D. melanogaster* and *C. elegans*. We report here a comprehensive study of the presence and absence of DNA methyltransferases (DNMTs) in 138 Ecdysozoa, covering Arthropoda, Nematoda, Priapulida, Onychophora, and Tardigrada. Three of these phyla have not been investigated for the presence of DNA methylation before. We observe that the loss of individual DNMTs independently occurred multiple times across ecdysozoan phyla. We computationally predict the presence of DNA methylation based on CpG rates in coding sequences using an implementation of Gaussian Mixture Modelling, `MethMod`. Integrating both analysis we predict two previously unknown losses of DNA methylation in Ecdysozoa, one within Chelicerata (Mesostigmata) and one in Tardigrada. In the early-branching Ecdysozoa *Priapulus caudatus* we predict the presence of a full set of DNMTs and the presence of DNA methylation. We are therefore showing a very diverse and independent evolution of DNA methylation in different ecdysozoan phyla spanning a phylogenetic range of more than 700 million years.

## Introduction

DNA methylation is prominent in vertebrates, where it is considered a fundamental part of epigenetic programming Lyko (2018). In human, about 70-80% of CpGs are methylated. Several non-vertebrate model organisms, such as *Drosophila melanogaster, Caenorhabditis elegans* and *Saccharomyces cerevisiae* Zemach et al. (2010); Raddatz et al. (2013) lack DNA methylation. It was discovered early-on, however, that some insects must have a DNA methylation mechanism Devajyothi and Brahmachari (1992). Since then, several studies have investigated the heterogenous distribution of DNA methylation in insects Field et al. (2004); Bewick et al. (2017); Provataris et al. (2018) and other arthropods de Mendoza et al. (2019b); Gatzmann et al. (2018). These showed that most insect orders have kept some amount of DNA methylation. The most prominent counterexample are Diptera which include the genus Drosophila. In nematodes, DNA methylation has only been identified in a few species. The highest levels are found in *Romanomermis cuicivorax* and low amounts in *Trichinella spiralis, Trichuris muris* and *Plectus sambesii* Gao et al. (2012); Rosié et al. (2018) suggesting an early loss during nematode evolution, prior to the separation of the nematod clades III, IV, and V. In most non-bilaterian metazoans DNA methylation is present, with the exception of placozoans de Mendoza et al. (2019a); Xu et al. (2019).

DNA methylation is a crucial mechanism in vertebrate gene regulation that plays a major role in cell fate decision making but their role in invertebrate gene regulation is much less clear. It appears that its function might differ significantly in different invertebrate groups. In the last years several experimental methods for detecting genomic DNA methylation have been developed. Nevertheless, they are still more expensive compared to sequencing the unmodified genome only. This can be problematic if one wants to widen the phylogenetic range of DNA methylation studies and include a large number of species. Another problem is that some of the lesser studied taxa are difficult to collect and culture which makes them less available for extensive experimental work. Bioinformatic studies such as the present one can help design such experimental studies. Relying on available public data we can make detailed predictions about the presence or absence of DNA methylation and the respective enzymes. Using these computational results one can decide more efficiently which taxa are most valuable to study to gain a new insight into the evolution of DNA methylation in invertebrates.

In animals, DNA methylation predominantly occurs at CG sites Goll and Bestor (2005); Lyko (2018). Two different sub-classes of enzymes are responsible for establishing DNA methylation. DNA methyltransferase 1 (DNMT1) reestablishes methylation on both DNA strands after a cell division. It preferentially targets hemi-methylated site. DNA methyltransferase 3 (DNMT3) can perform *de novo* methylation of unmethylated CpGs in the DNA. In vertebrates, DNMT3 is mainly active during embryonic development. However, the view of a clear separation of tasks has has been challenged Jeltsch and Jurkowska (2014); Lyko (2018). Not only does DNMT3 contribute to the maintenance of DNA methylation, DNMT1 has a notable *de novo* activity, as well. In addition DNMT1 might have other functions outside of DNA methylation Yarychkivska et al. (2018); Schulz et al. (2018) but they have not been studied extensively. Other functions are difficult to investigate, mainly because DNMT1 or DNMT3 knock-outs in human embryonic stem cells or mouse embryos have catastrophic consequences, e.g. cell death or embryonic lethality Liao et al. (2015).

DNMT2 has been believed to be a DNA methyltransferase as well until it was discovered that it recognizes tRNAs as a substrate. It methylates cytosine C38 of tRNA(Asp) in human and therefore is actually an RNA methyltransferase Goll et al. (2006).

DNA methyltransferases are believed to have emerged in bacterial systems from “ancient RNA-modifying enzymes” Iyer et al. (2011). Subsequently, six distinct clades of DNA methyltransferases have been acquired by eukaryotic organisms through independent lateral transfer Iyer et al. (2011). The DNMT clades thus do not have a common ancestor within the eukaryotes. DNMT1 and DNMT2 can be detected in most major eukaryotic groups, including animals, fungi and plants. Fungi lack DNMT3 but retained DNMT4 and DNMT5 similar to some, but not all, Chlorophyta (green algae). Embryophyta (land plants) lack DNMT4 and DNMT5 but harbor chromomethylase (Cmt), an additional DNA methytransferase related to DNMT1 Huff and Zilberman (2014). In Eumetazoa only DNMT1, DNMT2 and DNMT3 can be found. Although DNA methylation clearly is an ancestral process, it is not very well conserved among Protostomia.

All DNA methyltransferases (DNMTs) have a catalytic domain at their C-terminus. It transfers a methyl group from the substrate S-AdoMet to the C5 atom of an unmethylated cytosine Lyko (2018). However, the different families of DNMTs can be distinguished by their regulatory domains and conserved motifs in the catalytic domain Jurkowski and Jeltsch (2011). With five domains, DNMT1 has the most regulatory domains, see Figure 1 for an overview. The DMAP-binding domain binds DMAP1, a transcriptional corepressor. Also HDAC2, a histone deacethylase, establishes contact to the N-terminal region of DNMT1 Rountree et al. (2000). The RFTS domain (or RFD) targets the replication foci and directs DMAP1 and HDAC2 to the sites of DNA synthesis during S phase Rountree et al. (2000). The CXXC domain is a zinc-finger domain that can be found in several chromatin-associated proteins and binds to unnmethylated CpC dinucleotides Bestor (1992). The two BAH (bromo-adjacent homology) domains have been proposed to act as modules for protein-protein interaction Song et al. (2011); Yarychkivska et al. (2018).

**Fig. 1.**
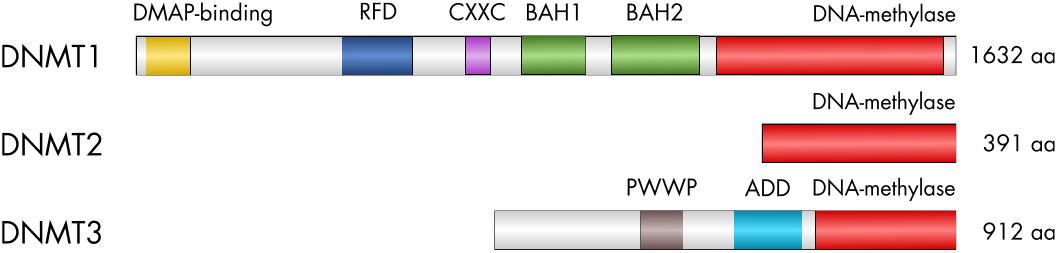
Conserved domains of animal DNA methyltransferases. Scaling and numbers refer to the human homologs.

DNMT3 has only two regulatory domains, a PWWP domain, named after the conserved Pro-Trp-Trp-Pro motif, and an ADD domain. Both mediate binding to chromatin. For the PWWP domain of (murine and human) DNMT3A, recognition of histone modifications H3K36me3 and recently also H3K36me2 has been reported Dhayalan et al. (2010); Weinberg et al. (2019). The ADD domain, is an atypical PHD finger domain, shared between ATRX, DNMT3, and DNMT3L, and has been shown to interact with histone H3 tails that are unmethylated at lysine 4 Zhang et al. (2010); Ooi et al. (2007).

DNMT2 has no regulatory domains Lyko (2018).

Methylated DNA is subject to spontaneous deamination of 5-methylcytosine, which leads to the formation of thymine and, consequently, to T·G mismatches. Over time, this results in C to T transition mutations predominantly in the context of CpG sites and CpG depletion in frequently methylated regions of the DNA. This changes the the number observed CpGs observed relative to the number expected from the C/G content of the genome. The observed/expected CpG distribution has been used in several studies to infer the presence of DNA methylation Bewick et al. (2017); Provataris et al. (2018); Aliaga et al. (2019); Thomas et al. (2020).

In *Apis mellifera* it has been show that its genes can be divided in two classes, depending on whether they exhibit a low or a high amount of CpG dinucleotides. This was explained by the depletion of CpG dinucleotides if DNA methylation is present. The highly methylated (low CpG) genes were associated with basic biological processes while lowly methylated (high CpG) genes were enriched with functions associated with developmental processes Elango et al. (2009). This “bimodal distribution” of CpG dinucleotides can be used to predict the presence of DNA methylation.

In invertebrates, gene bodies are methylated more heavily than other parts of the genome. Higher methylation levels should lead to a stronger statistical signal and therefore make it easier to decide if DNA methylation is present or not. Therefore, gene bodies have recently been in the focus of studies investigating DNA methylation in invertebrates. Several different criteria have been developed to distinguish the patterns of methylated and unmethylated DNA.

Bewick et al. (2017) use Gaussian mixture modeling (GMM) modeling with two components. Subsequently, they compare the 95% confidence intervals (CI) of the means. If they are overlapping they assumed a unimodal distribution, otherwise a bimodal one. In case of a bimodal distribution the presence of DNA methylation is assumed. Provataris et al. (2018) use the same GMM modelling. They define three different modes: “Bimodal depleted”, if the difference between both means is > 0.25 and the distribution with the lower O/E CpG ratio has a mean < 0.7, and the smaller component contains a proportion of the data > 0.1; “unimodal, indicative of DNA methylation”, if they do not fall in the first category but the portion of data which falls in the distribution with the lower O/E CpG ratio is ≥ 0.36 (this cutoff represents the corresponding value in *Bombyx mori*). All other cases are classified as “unimodal, not indicative of DNA methylation”. Aliaga et al. (2019) use a method based on kernel density estimations. They define four clusters based on the mode number (n), mean of the modes, skewness (sk) and standard deviation (sd). Three of the clusters are defined, among other parameters, as having one mode: “Ultra-low gene body methylation”, “Low gene body methylation” and “Gene body methylation”. Cluster with two modes (or 1 mode with skewness < −0.04) are defined as “Mosaic DNA methylation type”.

The predictions of the different methods are largely consistent although they may differ in individual cases and do not always match the the observed presence or absence of DNMTs, see section below.

In this paper, we present a detailed investigation of the presence and absence of DNA methyltransferases (DNMTs) across five ecdysozoan phyla, see Figure 2. Most of the 138 species analyzed here are from the phyla Arthropda and Nematoda. However, we also include less commonly studied groups such as Tardigrada, Onychophora and Priapulida. We identify at which points of the ecdysozoan evolution DNMTs were lost and investigate whethere there are common patterns between the phyla. In addition, we present an easy-to-use statistical approach for predicting the presence of genomic DNA methylation based on coding sequence data and apply it to our species of interest. The results of the predictions are compared with available experimental data.

**Fig. 2.**
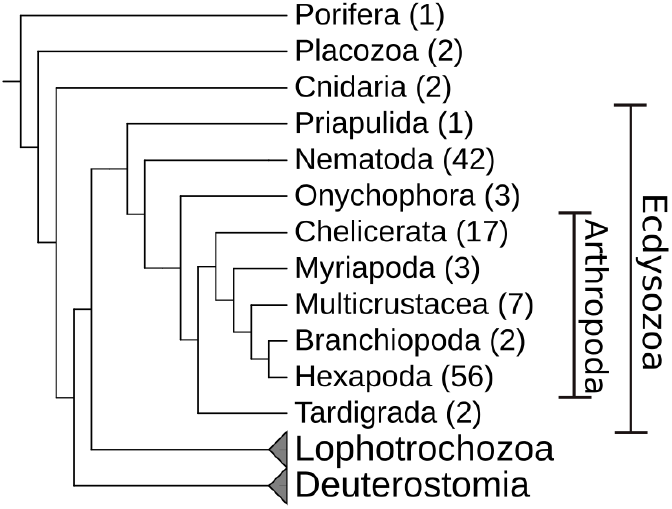
Overview of the metazoan phylogeny with a focus on Ecdysozoa. The number of species per group used in this study is given in brackets. Lophotrochozoa and Deuterostomia are shown for orientation only.

## Materials & Methods

### Identification of DNA methyltransferases

#### Proteome-based search

The predicted proteins of the species analyzed were downloaded from different sources, see supplementary Table 1. For 82 and 42 species data was taken from NCBI Sayers et al. (2019) and Wormbase Harris et al. (2020), respectively. Data for seven species each were retrieved from ENSEMBL Yates et al. (2020) and Laumer *et al*. Laumer et al. (2019).

The protein domain models for DNA_methylase (PF00145), ADD_DNMT3 (PF17980), CH (PF00307), PWWP (PF00855), BAH (PF01426), DMAP_binding (PF06464), DNMT1-RFD (PF12047) and zf-CXXC (PF02008) were downloaded from the “Pfam protein families database” El-Gebali et al. (2019). Initially, only the DNA_methylase model was used to identify DNA methyltransferase (DNMT) candidates in the set of proteins predicted using `hmmsearch` from the HMMER software http://hmmer.org/ version 3.2.1. Proteins with a predicted DNA_methylase domain and a full sequence e-value < 0.001 were further considered as candidates. For these, all before mentioned protein domains were annotated. Finally, each DNMT candidate was classified into one of three classes using custom perl scripts. A **DNMT1 candidate** was required not to have a PWWP or ADD_DNMT3 domain. In addition, having a DNMT1_RFD, zf-CXXC and BAH domain it was considered a *full* DNMT1 candidate, with only one of them a *partial* DNMT1 candidate. A **DNMT3 candidate** was required not to have a DNMT1_RFD, zf-CXXC or BAH domain. With both, a PWWP and a ADD_DNMT3 domain, it was considered a *full* DNMT3 candidate, with only one of them a *partial* DNMT3 candidate. A **DNMT2 candidate**, was required to have only a DNA_methylase domain and none of the other domains mentioned above. An overview of the required domains during the classification can be found in Supplementary Table 7. In a last step, the classification of the DNMT candidates was checked manually. The sequences of the DNA methylase domain of each candidate was extracted and aligned using `Clustal Omega` Sievers et al. (2011) version 1.2.4. A phylogenetic network was computed with *SplitStree4* Huson and Bryant (2006) version 4.10 and inspected manually for phylogenetic congruence of gene and species phylogeny. In case of contradicting results the specific conserved sequence motifs of the methylase domain were inspected manually and the candidate reassigned to a different class or discarded if it did not contain the proper sequence motifs Jurkowski and Jeltsch (2011).

#### Genome-based search

For selected subgroups an additional genome-based search for DNA methyltransferase (DNMT) candidates was performed. This was the case when the previously described workflow showed an unexpected absence of DNMTs in individual species. For example, a DNMT enzymes is detected in most species of a subgroup but is missing in one or two species. The groups that have been analyzed in addition were: Coleoptera for DNMT1 and DNMT3, Hymenoptera for DNMT3, Hemiptera for DNMT3, Chelicerate for all three DNMTs and Nematoda for all DNMTs. For each group, the DNMTs detected in the group, were used as queries. The programm `BLAT` Kent (2002) was used to search the query proteins against the species genome whenever the respective DNMT could not be found in the proteome. The script `pslScore.pl` (https://genome-source.gi.ucsc.edu/gitlist/kent.git/raw/master/src/utils/pslScore/pslScore.pl) available from the UCSC genome browser was used to assign a score to each genomic hit. The resulting bed-file was post-processed with the tools of the suite `bedtools` Quinlan and Hall (2010). All hits were clustered using `bedtools cluster`. If there were overlapping hits, only the best-scoring one was kept. Using blast-type output files from *BLAT* the genomic sequence to which the query was aligned could be extracted to get the full amino acid sequence corresponding to the hit. The full-length protein candidates were aligned using *Clustal Omega*. A phylogenetic network was computed with *SplitStree4* and inspected manually for phylogenetic congruence of gene and species phylogeny. Candidate proteins were discarded if they did not contain the methylase domain-specific, conserved sequence motifs. Otherwise they were kept as DNMT candidates.

This method allowed us to identify six additional DNMT enzymes in five species: *Asbolus verrucosus* DNMT1, *Soboliphyme baturini* DNMT2, *Acromyrmex echinatior* DNMT3, *Laodelphax striatellus* DNMT3, *Trichonephila clavipes* DNMT1 and DNMT3.

#### Inference of DNA methylation from CpG O/E value distributions

Coding sequences (CDS) for all species were downloaded from NCBI, Wormbase and ENSEMBL according to Supplementary Table 1. For the 7 species from *Laumer et al*. Laumer et al. (2019) this data was not available. We used two different datasets: the actual CDS data and shuffled CDS data. For the shuffled CDS data we performed a mononucleotide shuffling of the CDS data of each species using `MethMod`. The following analysis were performed for both the actual and the shuffled data.

For each CDS the Observed-Expected CpG ratio was calculated using the formula:

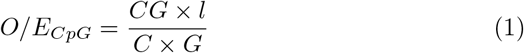

with *C, G*, and *CG* being the number of the respective mono- and dinucleotides in the given CDS and *l* being the length of the CDS. CDS shorter than 100 nucleotides or with more than 5% of N’s in the sequence were excluded.

We used a Gaussian Mixture Model (GMM) to identify possible subpopulations in the O/E CpG distribution. The Expectation Maximization algorithm in the python module ‘sklearn’ from the library *scikit-learn* Pedregosa et al. (2011) version 0.23.1 was used to estimate the parameters. The GMM was modeled with one or two components. For the GMM with one component, we calculated the Akaike information criterion (AIC). For the GMM with two components, we calculated the AIC and in addition the mean of each component, the distance *d* of the component means and the relative amount of data points in each component, see supplementary Table 2 and 3. For the distribution of O/E CpG values, the distribution mean, the sample standard deviation, and the skewness were calculated as well. All pairs of parameters were analyzed using two-dimensional scatterplots generated with R.

We used the distance between the component means as an indicator for DNA methylation. If the distance is greater or equal to 0.25, we assume DNA methylation is present, otherwise it is absent.

### Ecdysozoan Phylogeny

The topology of the ecdysozoan phylogeny, used for display only, is a composite of phylogenetic information compiled from several studies. The topology of Arthropoda was based on Misof et al. (2014) and combined with phylogentic information for the taxa Coleoptera Zhang et al. (2018), Lepidoptera Kawahara et al. (2019), Hymenoptera Peters et al. (2017), Hemiptera Johnson et al. (2018), Aphididae von Dohlen et al. (2006); Kim et al. (2011); Nováková et al. (2013), Crustacea Schwentner et al. (2017), Copepoda Khodami et al. (2017), Chelicerata Howard et al. (2020); Sharma et al. (2012), Aranea Fernández et al. (2018), and Acari Arribas et al. (2020). The topology of the nematode phylogeny was based on Consortium et al. (2019) and combined with phylogenetic information for the genera Plectus Rošić et al. (2018), Trichinella Korhonen et al. (2016), Caenorhabditis Stevens et al. (2019), and Diploscapter Fradin et al. (2017).

## Results

### Presence and absence of DNA methyltransferases in Ecdysozoa species

We investigated the presence of DNMTs in 138 species using a carefully designed homology search strategy (see Materials and Methods) aiming at minimizing false negatives. Candidate sequences were then curated carefully to avoid overprediction. Most of the available genomes belong to the Nematoda (42) and Arthropoda (85). Of the arthropod species, 56 are Hexapoda (insects) and 29 belong to other subphyla. Only 6 species are from Ecdysozoa groups outside of Nematoda or Arthropoda. In addition 5 species from groups outside of Bilateria have been included. In seven species, the arthropods *Calanus finmarchicus, Eudigraphis taiwaniensis, Glomeris marginata, Anoplodactylus insignis* and all three Onychophora species, no genome data was available but only proteins predicted from transcriptomic data. The respective species are indicated in the text by stating that they have a “transcriptome only” (t.o.).

Our findings are summarized in Figures 3, 4, 5 and supplementary Figure 1. Potential losses of DNMT1, DNMT2, and DNMT3 are marked with stars in the respective colors. Species with a transcriptome only (t.o.) are indicated by triangles. In the following paragraphs we discuss the results of our annotation efforts in more detail.

**Fig. 3.**
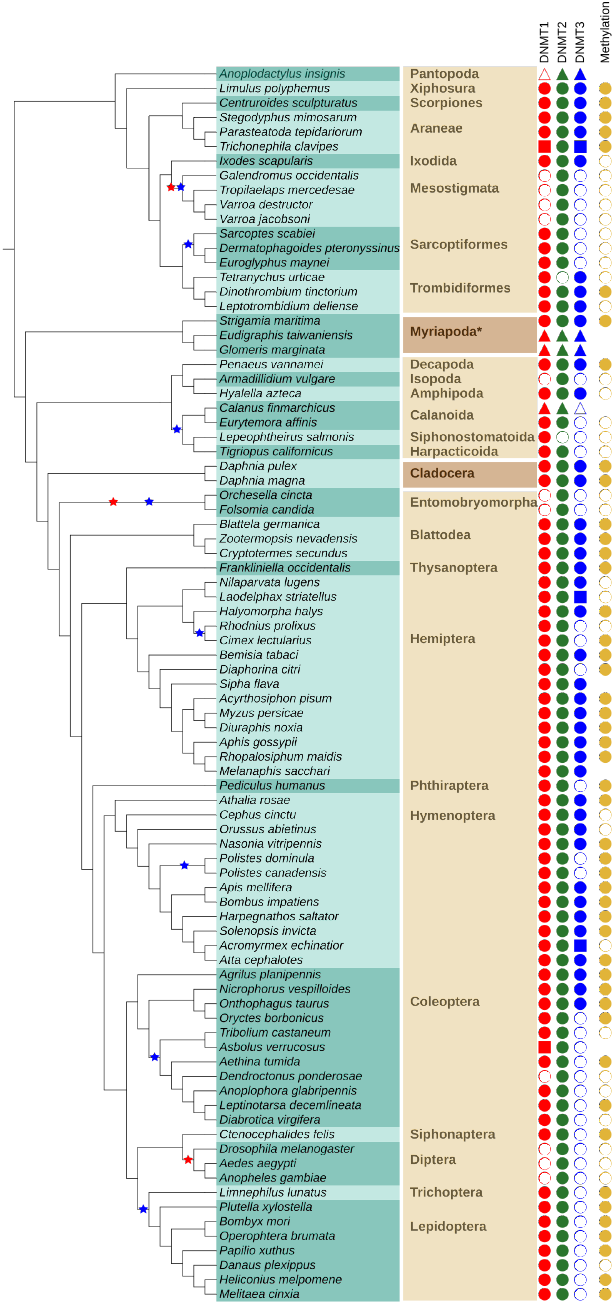
Presence and absence of DNMT family members in Arthropoda indicated by filled and open symbols, respectively for DNMT1 (red), DNMT2 (green), and DNMT3 (blue). Data sources are indicated by symbol shape: proteome ○, genome □, transcriptome △. The rightmost column (golden circles) shows the presence and absence of DNA methylation as predicted from the O/E CpG ratio. Absence of golden circle indicates missing data. The species list is given on turquoise background with alternating shades indicating the order membership. The name of the order (or suitable higher group marked with an asterisk *) is given in bold. Alternating shades of brown indicate (from top to bottom) Chelicerata, Myriapoda, Multicrustacea, Branchiopoda, and Hexapoda. Stars in the species tree denote proposed loss events inferred from absence of a DNMT in all species of a subtree comprising at least two leaves, disregarding absences in species with transcriptomic data only.

#### Arthropoda

Arthropoda are an extremely species-rich and frequently studied group of invertebrates. The most prominent subphylum is Hexapoda, which contains, among others, all insects. Several (emerging) model organism belong to insects, e.g. the fruit fly *Drosophila melanogaster* (Diptera), the silk moth *Bombyx mori* (Lepidoptera), the red flour beetle *Tribolium castaneum* (Coleoptera) or the honey bee *Apis mellifera* (Hymenoptera). The group of Crustacea (crabs, shrimp, lobster) is currently believed to be paraphyletic Schwentner et al. (2017). Multicrustacea consists of most of the “crustacean” species, e.g. the white leg shrimp *Penaeus vannamei* (Decapoda) or the amphipod *Hyalella azteca* (Amphipoda). Branchipoda with the frequently studied water flea *Daphnia pulex* (Cladocera) are currently placed more closely related to Hexapoda. The sister group to all of the aforementioned groups are Myriapoda (millipedes, centipedes). The earliest-branching group of Arthropoda are the Chelicerate. A diverse subgroup of Chelicerata are Arachnida (e.g. spiders, scorpions, ticks) but they also contain the Atlantic horseshoe crab *Limulus polyphemus* (Xiphosura) and sea spiders (Pantopoda). We analyzed 85 species of the phylum Arthropoda. They belong to 28 different taxonomic orders. An overview of the results can be found in Figure 3.

The subphylum Hexapoda was the largest group analyzed with 11 different orders. Two had a full set of DNMTs: Blattodea (3 species) and Thysanoptera (1). In four orders only DNMT1 and DNMT2 are present: Siphonaptera (1), Trichoptera (1), Lepidoptera (8) and Phthiraptera (1). In two only DNMT2 could be identified: Diptera (3) and Entomobryomorpha (2). In the remaining three orders the occurrence of DNMT enzymes is heterogeneous suggesting secondary losses within the order. Coleopetera (11 species) have all DNMTs, DNMT1 and DNMT2 or only DNMT2. Hymenoptera (12) mostly have all DNMTs but in two species of the genus Polistes, DNMT3 could not be detected. In three species of Hemiptera (14) we did not find DNMT3, as well.

The subphylum Crustacea is currently believed to be paraphyletic Schwentner et al. (2017) but the following species are considered part of it. In two species of the Daphnia genus all DNMTs have been found. They belong to the order Cladocera in the class Branchiopoda, formerly part of the subphylum Crustacea. Six additional orders of the former subphylum, belonging to the group of Multicrustacea have been studied. In Amphipoda (1) and Decapoda (1) all three DNMTs have been found, as well. In the orders Calanoida (2 species), Harpacticoida (1) and Siphonostomatoida (1) DNMT3 was not identified. In the calanoida *Lepeophtheirus salmonis* DNMT2 could not be identified as well. In Isopoda (1) DNMT1 and DNMT3 could not be detected.

In the subphylum Myriapoda three different orders have been analyzed with one species each. All of them showed a full set of three DNMT enzymes.

17 species of the subphylum Chelicerata were analyzed. They belong to 8 different orders. We detected all three DNMTs in Xiphosura (1 species), Scorpiones (1), Aranea (3) and Ixodida (1). The same was the case for Trombidiformes (3) with the exception of *Tetranychus urticae* for which DNMT2 could not be found. In Sarcoptiformes (3) only DNMT3 was not detectable. In Mesostigmata (4) this was the case for DNMT1 and DNMT3. In the one species of Pantopoda (1) *Anoplodactylus insignis* (t.o.) DNMT1 could not be found.

#### Nematoda

Nematoda are, next to Arthropoda, the best-studied group of Ecdysozoa. Developing a complete nematode systematics is still an ongoing process.

Most available genome data comes from the clades I, III, IV and V. Clade V contains the most well-known nematod species *Caenorhabditis elegans*.

42 nematodes species of five clades were analyzed. Of the 17 species in clade V most had no DNMTs, in 5 species DNMT2 could be detected. In clade III for 8 out of 10 species DNMT2 was present but not the other DNMTs. Clade IV with six species showed no signs of DNMT at all. In *Plectus sambesii*, the only representative of its clade, DNMT3 could not be found. In clade I, in 6 of the 8 species only DNMT2 and DNMT3 were detected. For one species all three DNMTs have been identified. In another one species only DNMT3 is present but DNMT2 could not be found. An overview of the results can be found in Figure 4.

**Fig. 4.**
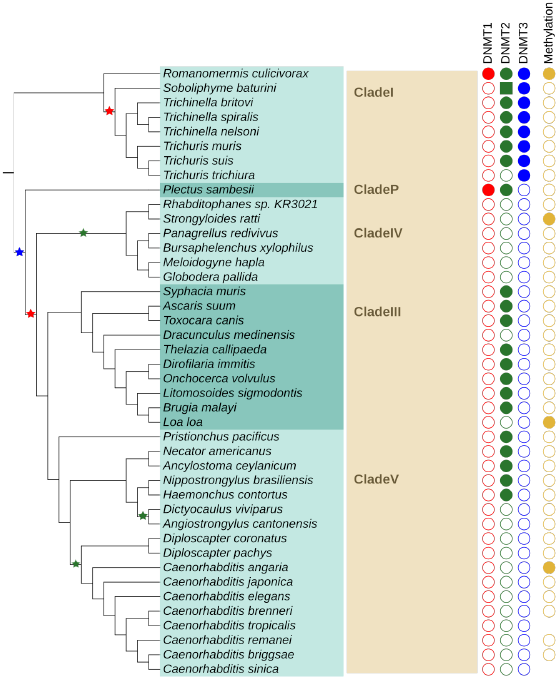
Presence and absence of DNMT family members in Nematoda. See Fig. 3 for detailed legend. Instead of order names, clade names are given (in bold).

#### Priapulida, Onychophora and Tardigrada

These groups are not often in the focus of scientific studies. Tardigrada, commonly known as water bears, gained some interest because they can survive in very harsh conditions, such as extreme temperature, radiation, pressure, dehydration and even in outer space Jönsson et al. (2008). Onychophora or velvet worms are the sister taxon to Arthropoda+Tardigrada. Some species can bear live offsprings Ostrovsky et al. (2016). Priapulida (penis worms) are believed to be among the earliest branching Ecdysozoa and therefore are of great interest for comparative studies. Unfortunately, genomic data so far is only available for one species.

In the Onychophora (3) (t.o.) *Peripatoides sp*. and *Peripatopsis overbergiensis* DNMT1 and DNMT2 was detected in *Peripatus sp*. DNMT2 and DNMT3. In Tardigrada (2) only DNMT2 could be identified. In the single member of the Priapulida all DNMTs were detected. An overview of the results can be found in Figure 5.

**Fig. 5.**
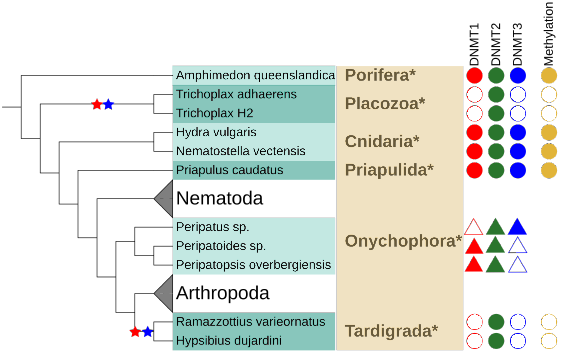
Presence and absence of DNMT family members in Priapulida, Onychophora and Tardigrada and early branching Metazoa. See Fig. 3 for detailed legend.

#### Early-branching Metazoa

The systematics of early-branching Metazoa is difficult to resolve and currently still heavily discussed. The Cnidaria (jellyfish, sea anemones, corals) are believed to be the closest relatives to bilateral animals. Placozoa are a more distant taxa with Trichoplax as the most prominent genus. They are tiny and delicate marine animals. For a long time only one species *Trichoplax adhaerens* was known along with a number of haplotypes. Only recently two more species have been described. Porifera, or sponges, are (together with Ctenophora) a contender for being the earliest branching phylum of Metazoa. In the outgroup Placozoa (2) only DNMT2 was detected while in Cnidaria (2) and Porifera (1) all DNMT enzymes were found.

### DNA methylation inferred from CpG O/E value distributions

The ratio of observed and expected CpGs serves as an indicator for the presence of DNA methylation. In invertebrates often only a subset of genes is subject to CpG methylation. Therefore, we assume that the observed distribution is a mixture of two gaussian distributions. Similar to previous work, we use an expectation–maximization (EM) algorithm to estimate the parameters of this Gaussian Mixture Model (GMM) Bewick et al. (2017); Provataris et al. (2018). The results outlined below were used to revise the parameters reliably indicating bimodality and thus the presence of DNA methylation.

Coding sequence (CDS) data was available for all species except *Calanus finmarchicus, Glomeris marginata, Eudigraphis taiwaniensis, Anoplodactylus insignis*, *Peripatopsis overbergiensis*, *Peripatoides sp*., *Peripatus sp*., whose data was from *Laumer et al*. Laumer et al. (2019). For five species *(C. sinica, C. tropicalis, S. flava, M. sacchari, A. verrucosus*) the genome was not published, yet, therefore they have been excluded from this genome-wide analysis. Hence we were able to analyze O/E CpG ratios for the CDS of 126 species.

We performed Gaussian Mixture Modelling (GMM) with the actual CDS data and a mononucleotide shuffled version of the CDS data. The later served as a negative control since CpG dinucleotide depletion is not to be expected. To evaluate whether a model with one or two components better represents the observed CpG O/E distribution, we first applied the Akaike Information Criterion (AIC), which is a measure of relative goodness of fit. For 94 of 128 species a model with two components was favored over a model with one component. In contrary to our expectation, the AIC also favored a two component model for 94 of 128 shuffled CDS data.

This indicates that the CDS may also fall into two classes distinguished by overall GC content, not only by relative CpG abundance. Although the AIC is generally accepted for GMMs, empirically, we find that the AIC is a poor decision criterion for our purposes.

Features directly derived from the two components, such as the component means and the relative amount of data points corresponding to each component clearly proved to better separate real and shuffled data. Table 1 shows that the mean distance between the two components is much larger in the real data compared to the shuffled data. Hence we use the difference between the means of the two Gaussians as an indicator of CpG depletion. As the distance is continuous, ranging from 0.00 to 0.63 in our data, it is necessary to determine the threshold above which the difference of two means is interpreted as indicative of DNA methylation. Naively, species having neither DNMT1 or DNMT3 should be less likely to contain DNA methylation, while species in which one or both of the enzymes are present should be more likely to have kept genomic DNA methylation. Of the 126 species analyzed, in 45 the DNMT1 and DNMT3 enzymes have been found, while in 46 neither was found. In 28 species only DNMT1 was detected and in 7 species only DNMT3, see Table 2. Figure 6 shows the means of both GMM components for all analyzed species, marked by different colors and symbols according to their set of DNMT1/3 enzymes and their taxonomic group. The threshold value *d* ≥ 0.25 is able to separate almost all of the species with no DNMT1/3 from the others. We have chosen this conservative threshold in order to avoid false positive prediction of DNA methylation. In our data, 55 of 126 species had a distance greater or equal to 0.25 indicative of DNA methylation. The other 71 species had a distance smaller than 0.25.

**Fig. 6.**
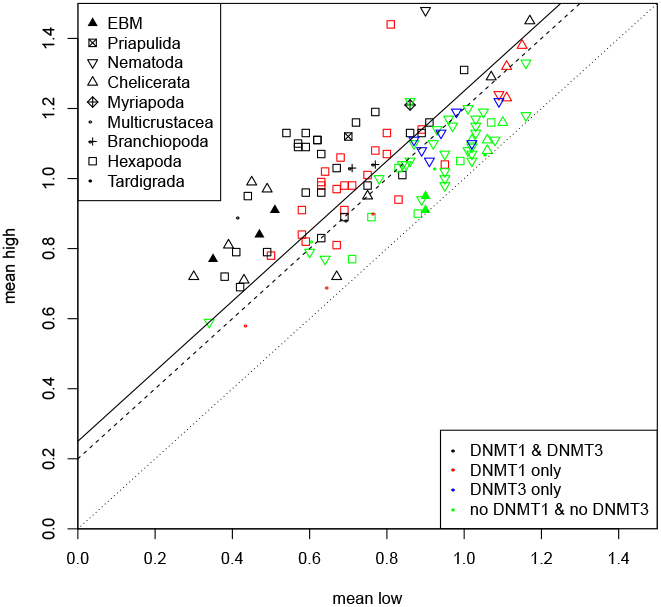
Each point shows one species analyzed by Gaussian Mixture Modelling (GMM). The axes are the means of the two components. The taxonomic group is indicated by the style of the point. The color represents if both, DNMT1 and DNMT3 (green), have been found in the species, only DNMT1 (red), only DNMT3 (black) or neither one nor the other (blue). The diagonal lines indicate the distance between the mean of both GMM components. The dotted line indicates a distance of *d* = 0, the dashed one *d* = 0.2 and the solid line *d* = 0.25 (selected threshold). ‘EBM’ stands for ‘Early-branching metazoa’, i.e. Porifera, Placozoa and Cnidaria.

**Table 1.** Summary of the Gaussian Mixture Modelling for real and shuffled data. “mean-Low” and “meanHigh” are the component means corresponding to the components with lower and higher O/E CpG ratios (first and second row). The distance *d* between the means is given in the third row. “%low” gives the relative amount of data points (transcripts) in the component with the lower O/E CpG ratio, “%low” + “%high” equals to 1. Due to its extreme values the nematode *Loa loa* was excluded from this table. Its values are: “mean-Low” 1/1, “meanHigh” 4.53/1.18, d 3.55/0.18 and “%low” 0.99/0.98 for the real/shuffled data.

| Arthropoda |  |  |  |  |  |  |
| --- | --- | --- | --- | --- | --- | --- |
| Range | Real data |  |  | Shuffled data |  |  |
|  | Min. | Mean | Max. | Min. | Mean | Max. |
| meanLow | 0.30 | 0.72 | 1.17 | 0.95 | 0.99 | 1.00 |
| meanHigh | 0.58 | 1.00 | 1.46 | 1.00 | 1.02 | 1.05 |
| distance $d$ | 0.01 | 0.28 | 0.63 | 0.00 | 0.03 | 0.11 |
| %low | 0.14 | 0.46 | 0.87 | 0.37 | 0.72 | 0.81 |
| Nematoda |  |  |  |  |  |  |
| Range | Real data |  |  | Shuffled data |  |  |
|  | Min. | Mean | Max. | Min. | Mean | Max. |
| meanLow | 0.34 | 0.94 | 1.16 | 0.93 | 0.98 | 1.00 |
| meanHigh | 0.59 | 1.10 | 1.48 | 1.00 | 1.02 | 1.07 |
| distance $d$ | 0.00 | 0.15 | 0.58 | 0.00 | 0.04 | 0.14 |
| %low | 0.13 | 0.59 | 0.96 | 0.49 | 0.74 | 0.82 |

**Table 2.** Relationship between the combination of DNMT candidates and the predicted methylation level. Shown is the amount of species for which DNA methylation is predicted to be present or absent classified by the presence of DNMT enzyme combinations.

| enzymes present | total | methylation |  |
| --- | --- | --- | --- |
| | | present<br>$d \geq 0.25$ | absent<br>$d < 0.25$ |
| DNMT1 & DNMT3 | 45 | **36** | 9 |
| DNMT1 only | 28 | 16 | 12 |
| DNMT3 only | 7 | 0 | 7 |
| no DNMT1 & no DNMT3 | 46 | 3 | **43** |
|  | 126 | 55 | 71 |

## Discussion

To our knowledge this study is the phylogenetically most diverse analysis of DNA methylation in Ecdysozoa, to-date. Several recent projects have investigated DNA methylation in species of Ecdysozoa but they have focused on different subgropus, i.e. Hexapoda Bewick et al. (2017); Provataris et al. (2018), Arthropoda Lewis et al. (2020); Thomas et al. (2020) and Nematoda Rošić et al. (2018). In our study we have investigated a similar number of orders from the before mentioned groups. Of the arthropod subphylum Chelicerata we included a larger number of orders and therefore were able to predict an additional loss of DNA methylation. In addition we included species from Priapulida, Onychophora, and Tardigrada. The presence of DNA methylation has been investigated in none of these phyla before. In Tardigrada we predict an additional, previously unknown, loss of DNA methylation. All of our data was analyzed with the same computational pipeline for detecting DNA methyltransferase enzymes and predicting DNA methylation based on the CpG ratios. The results are therefore comparable over a large phylogenetic range, spanning more than 700 million years Kumar et al. (2017) of ecdysozoan evolution. Our analysis of five out of seven Ecdysozoa phyla confirms that the evolution of DNA methylation in Ecdysozoa proceeds independently in each phylum. It is therefore of great interest to perform experimental studies in each of these phyla to discover different evolutionary adaptations DNA methylation might have undergone.

### Presence and Absence of DNA methyltransferases

Overall, our data shows that both individual DNMTs and DNA methylation as a process have been lost independently in multiple lineages. Since the absence of an enzyme is difficult to prove conclusively, we rely on data from related species and invoke parsimonious patterns to identify loss events with confidence: the lack of evidence for a DNMT in an entire clade of related species makes a loss event a very plausible explanation.

There are several reasons why a DNMT may escape detection. The most prominent cause is a low quality, fragmented genome assembly. Not finding a homolog in a species with a high quality, completed genome assembly, in particular in model organisms such as *Caenorhabditis elegans* and *Drosophila melanogaster* makes a negative search result more reliable. It is also possible that a protein has diverged so far that it is no longer recognizable as a homolog in the target organism by the search method used. This explanation becomes more likely in groups, such as Tardigrada or Nematoda, where the closest known homolog of DNMT enzymes is quite far away. If they have diverged extensively it is more likely to miss existing DNMTs. Nevertheless, as long as the catalytic domain of the enzymes still performs the same function we should be able to find the enzyme. The predicted phyletic pattern of DNMT losses is quite different in Arthropoda and Nematoda. DNMT1 is found in most arthropod species analyzed in our study. Three independent loss events of DNMT1 are suggested by our data (3). In Nematoda only two events of DNMT1 loss are suggested but they occur earlier in the evolution of the studied nematod species. Therefore, only in two species DNMT1 can still be detected.

DNMT2 is most likely present in all Arthropoda. The absence in two individual species is probably a technical artifact since DNMT2 enzymes are present in closely related species in both cases. In Nematoda, absence of DNMT2 enzymes is fare more frequent. Given the near perfect conservation of DNMT2 in other metazoan species, this is rather unexpected. Interestingly, the candidate DNMT2 sequences are clearly more divergent compared to those in Arthropoda, which may hint at false positive predictions of 13 DNMT2 enzymes. In this case, a single loss event either after divergence of clade I or both, clade I and clade P, is plausible.

DNMT3 seems to be the most dispensable member of the DNMT family. According to our data, it was lost eight times in Arthropoda. It only occurs in combination with DNMT1 and is lost prior to or simultaneously with loss of DNMT1. In Nematoda, DNMT3 is present in all members of clade I and absent in all other clades. Interestingly, in all but one species of clade I, we detected a DNMT3 in the absence of DNMT1.

Absence of DNMT3 in the presence of DNMT1 is frequently associated with low levels of CpG depletion. The weak bimodality of the CpG ratio distribution may be the consequence of a return to an unbiased, unimodal distribution caused by decaying methylation levels due to failure to (re-)establish and maintain methylation. Under certain conditions, DNMT1 may have weak *de novo* activity Dahlet et al. (2020). The molecular mechanism involves binding to unmethylated CpGs via the CXXC domain and auto-inhibition of de novo methylation Song et al. (2011). Via its regulatory domains DNMT1 interacts with epigenetic factors which may be involved in regulalting DNMT1 *de novo* activity.

The loss events as defined in this study are well supported by the absence of the enzymes in related species, see the colored stars in Figures 3, 4 and supplementary Figure 1. More precisely, a loss is only inferred if the respective DNMT could not be found in all species of the respective subtree and if it contains at least 2 species. Considering the problems in gene detection, these rules remove cases where the poor quality of single genomes may prevent the detection of DNMTs. In Arthropoda all members of the DNMT family can be identified in several species of each subphylum. Therefore it is unlikely that the negative predictions are caused by extreme divergence of protein sequences that might have rendered them undetectable by homology search methods. The N50 value (that is, 50% of the genome is covered by contigs with a length of at least N50) serves a good measure of assembly quality for our purposes. In Arthropoda, five species are missing DNMT1 or DNMT3 and are not covered by the loss events we propose. The genomes of *Diaphorina citri* (Hemiptera), *Armadillidium vulgare* (Multicrustacea) and *Oryctes borbonicus* (Coleoptera) are the 13th, 8th and 7th worst assemblies in Arthropoda according to the N50 value, see supplementary Table 1. The N50 for *D. ponderosae* (Coleoptera) is around average and for *Anoplodactylus insignis* (Chelicerata) only a transcriptome is available. It is difficult therefore, to interpret these potential loss events. A more reliable prediction will be possible when better genomes or data from more closely related species becomes available.

The DNMT1/DNMT3 losses in Nematoda are more difficult to evaluate since there are so few positive findings. Their absence in clade III, IV and V is supported by the findings of Rošić et al. (2018). These groups contain several high quality genomes, such as the model organism *C. elegans*. The most likely reason for missing existing proteins would therefore be that they are already too diverged. However, DNA methylation has been verified to be absent in several of them and no findings of DNMT enzymes have ever been reported. Therefore, it seems reasonable to conclude that DNA methylation and both DNA metyltransferases are absent fron Nematoda of clade III, IV, and V.

In clade I, DNMT3 is evidently present. However, it seems that DNMT1 is absent in all but a single species examined. This pattern cannot be seen in any other ecdysozoan group. The exception is the earliest branching nematode *Romanomermis cuicivorax*, which posesses both, DNMT1 and DNMT3, as well as DNMT2. The case of *Plectus sambesii*, the sole member of clade P, is quite interesting because DNMT1 is present while DNMT3 is absent. However, the genome of *P. sambesii* is the 3rd worst of all nematods putting the loss of DNMT3 into question. We can therefore suggest two possible scenarios, either DNMT3 was lost in the stem lineage of clade P and the clades III, IV and V, i.e. before the loss of DNMT1 or after branching of clade P and clades III, IV and V and simultaenously with the loss of DNMT1.

The two missing DNMT2 in Arthropoda are likely to to be false negatives since homologs of DNMT2 were detected in all other arthropods. Likely, this is also the case in the nematode *Trichuris trichiura* since in the two other species of its genus DNMT2 was found. In clade III, IV, and IV the pattern is not very parsimonious and our analysis reports three independent DNMT2 loss events. In addition, we did not detect DNMT2 candidates in two more species in clade III. Visual inspection of the DNMT2 alignment revealed that DNMT2 candidates of clades III and V are highly divergent. In conclusion, it remains questionable whether these enzymes are still functional DNA methyltransferases.

Supplementary Tables 4, 5 and 6 summarize our results and provide a comparison with five recent studies. We analyzed 138 species in total, of which 37 and 34 have been previously examined by Bewick et al. (2017) and Provataris et al. (2018), respectively. The evolutionary history of DNMT1 within Hymenoptera, including paralogization, is described in more detail in Bewick et al. (2017) compared to our work. We have focused on determining if at least one copy of DNMT1/2/3 is present in a genome since we wanted to mainly study losses of DNA methylation in Ecdysozoa. To the largest part, the results of all studies are in concordance. We were able to identify DNMTs in seven species, i.e DNMT1 in two species (*P. vannamei* and *N. nevadensis*) and DNMT3 candidates in five species (*P. vannamei*, *I. scapularis*, *B. germanica, N. lugens* and *H. halys*), respectively, which have been missed in at least one other study. We, on the other hand, miss no DNMT enzyme reported by Bewick et al. (2017) or Provataris et al. (2018). Two subsequent studies de Mendoza et al. (2019a) and Lewis et al. (2020) have analyzed fewer Hexapoda but included other arthropods and some non-bilaterian species. We share 16 and 20 species with these studies. The results for detecting DNMTs are almost identical we find DNMT1 in one less species, *A. vulgare*, but DNMT3 in one more, *I. scapularis*, compared to Lewis et al. (2020). Of the 42 Nematoda analyzed in our study, Rošić et al. (2018) investigated a subset of 14. The results for the presence/absence of DNMT enzymes in these 14 species are identical.

### DNA methylation inferred from CpG O/E value distributions

Traditionally, a computational prediction for the presence of DNA methylation is considered to be much weaker evidence than an experimental verification, e.g. by bisulfite sequencing. In principle we agree that an experimental verification leads to a better insight about the actual distribution of DNA methylation in a genome. Nevertheless, aside from the additional work required to gain genomic DNA for each species and perform the experiments, there are fundamental differences between the results of experiments and our prediction. The results of bisulfite sequencing are specific for the tissue which was used to extract the genomic DNA, e.g. whole organims, body parts or particular developmental stages. Strictly speaking the results are only valid for the analyzed tissue. With our method of predicting the DNA methylation from the O/E CpG rates we basically analyze the DNA methylation in the germline. Only mutations (caused by deamination) which happen in the germline will be kept in the next generations. DNA methylation of germ cells is rarely measured experimentally in invertebrates due to the additional difficulties collecting enough genomic material. Therefore, contrary to most experimental approaches, we actually predict germline DNA methylation.

Over evolutionary time, the distribution of CpG dinucleotides is influenced by DNA methylation, which gives rise to an increased rate of C to T mutations and, consequently, CpG depletion. In case of genome-wide DNA methylation, as in vertebrates, the signal is easy to detect. The situation is more challenging in invertebrates, where methylation is often concentrated to a subset of coding regions. A two-component Gaussian Mixture modelling (GMM) approach is used to model the populations of methylated and unmethylated coding sequences. As we could show, the distance *d* between the component means is a reasonable measure for the level of DNA methylation in Ecdysozoa. Using *d* ≥ 0.25 as threshold we could confirm the previously reported *absence* of notable DNA methylation in several species, such as the fruit fly *Drosophila melanogaster* (*d* = 0.01), the red flour beetle *Tribolium castaneum* (*d* = 0.08) or the nematode *Caenorhabditis elegans* (*d* = 0.20). Furthermore, we predicted the *presence* of DNA methylation in a number of species such as, the insects *Bombyx mori* (*d* = 0.39), *Nicrophorus vespilloides* (*d* = 0.37), *Apis mellifera* (*d* = 0.58), *Acyrthosiphon pisum* (*d* = 0.49), *Blatella germanica* (*d* = 0.30), the water flea *Daphnia pulex* (*d* = 0.32) or the nematod *Romanomermis culicivorax* (*d* = 0.58), which is in concordance with the literature.

Unfortunately, the number of studies which used experimental methods to verify the presence of DNA methylation in Ecdysozoa is quite limited, in particular outside of Hexapoda. Our data suggests several loses of DNA methylation which can not be supported by evidence other than the computationally calculated O/E CpG ratio. Due to the predicted presence of DNA methylation in closely related species some “species-specific” losses seem questionable, e.g. *Danaus plexippus* (*d* = 0.11) and *Acromyrmex echinatior* (*d* = 0.24). Conversely, some of the positive findings are likely to be false predictions, e.g. the nematods *Caenorhabditis angaria* (*d* = 0.36), *Loa loa* (*d* = 3.55) and *Strongyloides ratti* (*d* = 0.25). For many other species there is currently no experimental verification available. The reason for the incorrect predictions is currently not easy to explain. Most likely, there are other, presently unknown factors that influence the distribution in CpGs in the genome. Such effects are difficult to distinguish from the effects of DNA methylation.

Nine species in which we detected DNMT1 and DNMT3 were predicted to not have DNA methylation. The chelicerata *I. scapularis* (*d* = 0.20), *T. urticae* (*d* = 0.06) and *L. deliense* (*d* = 0.22), the amphipod *H. azteca* (*d* = 0.18), the hemiptera *N. lugens* (*d* = 0.20) and *L. striatellus* (*d* = 0.20) and the hymenoptera *C. cinctu* (*d* = 0.22), *O. abietinus* (*d* = 0.17), *A. echinatior* (*d* =0.23).

*Tribolium castaneum* is one example where DNMT1 was kept despite the loss of DNA methylation Schulz et al. (2018) but there is currently no example known where both DNA methyltranserases (DNMTs) are kept despite the loss of DNA methylation. Therefore, one would assume species with both DNA methyltranserases are likely to have DNA methylation as well. Nevertheless, only for one of the nine species, *I. scapularis*, this was experimentally verified. It is likely that most of these cases are false negatives but without additional information one can not be sure. In species closely related to the chelicerata *T. urticae* and *L. deliense* we detected several losses of DNMTs as well as DNA methylation. The situation is similar for the the hemiptera *N. lugens* and *L. striatellus*. It is possible that DNA methylation has been significantly reduced in these groups and therefore can not be detected by our prediction method anymore.

Computational predictions of methylation status have been performed with different methods by *Bewick et al*. Bewick et al. (2017) and *Provataris et al*. Provataris et al. (2018). Supplementary Table 5 provides a summary of their findings and the respective results from our study. Compared to Bewick et al. (2017) there are three cases where we predict no DNA methylation while they predict DNA methylation: *R. prolixus* (*d* = 0.14), *O. abietinus* (*d* = 0.17) and *A. glabripennis* (*d* = 0.21). In one case *M. cinxia* (*d* = 0.27) we predict DNA methylation while they do not. Compared to Provataris et al. (2018), there are five cases where we predict DNA methylation while they do not: *S. maritima* (*d* = 0.35), *H. saltator* (*d* = 0.44), *A. cephalotes* (*d* = 0.27), *P. xylostella* (*d* = 0.28) and *M. cinxia* (*d* = 0.27). In one case, *D. plexippus*, they predict DNA methylation while we do not.

In total these are 9 species in which our methylation predictions disagree with at least one of the other two papers. In the case of *S. maritima* and *H. saltator* there is experimental evidence for DNA methylation so our prediction is backed up by that. For the other species no such data is available. The prediction of the presence of DNA methylation in *M. cinxia* is the only case where both other studies agree on contradicting our prediction. This species would be the only exception in Lepidoptera without DNA methylation, therefore our prediction appears to be more likely. In *A. glabripennis* we predict no DNA methylation while Bewick et al. (2017) does but there is no further evidence available. The other 5 species are part of all three studies and in all cases our prediction is supported (three times Bewick et al. (2017), two times Provataris et al. (2018)) by one study and contradicted by the other. There is no case where our predictions is clearly worse than those of competing methods. In the single case of *A. glabripennis* there is no further evidence to resolve a contradicting result.

For 28 of the species examined, experimental data on the presence (22) and absence (6) of DNA methylation is available. We correctly predict the presence and absence of DNA methylation for 17 and 6 species, respectively, totaling to 23 out of 28. The remaining five predictions are false negatives. Note that there are no false positive predictions given the experimental data set at hand. Among the species corresponding to the false negative predictions are two arthropod species, *I. scapularis* (*d* = 0.2) and *A. vulgare* (*d* = 0.21), and three nematode species *T. spiralis* (*d* = 0.24), *T. muris* (*d* = 0.08) and *P. sambesi* (*d* = 0.15), see also supplementary Table 4 and 5. According to Lewis et al. (2020), the level of DNA methylation in *A. vulgare* is very low which is likely the reason why our prediction method fails. There is no obvious explanation why we miss DNA methylation in *I. scapularis*. In the three nematodes, notable levels of DNA methylation are mostly present at repeats, which cannot be captured by our method. According to Rošić et al. (2018) only the nematod *R. culicivorax* shows a bimodal distribution for DNA methylation across genes.

### Conclusions

The amount of genomic and transcriptomic data from a wide range of species is constantly increasing. Often only a relatively small phylogenetic range is analyzed simultaneously. The analysis of “universal” evolutionary patterns, however, requires that the same analysis is applied to widely different groups of species. With this study we provide the most diverse analysis of DNA methyltransferase enzymes in Ecdysozoa, to date, spanning a phylogenetic range of more than 700 million years. Previous studies have focused on specific subgroups in particular Arthropoda Bewick et al. (2017); Provataris et al. (2018); Lewis et al. (2020); Thomas et al. (2020) and Nematoda Rošić et al. (2018) and covered only selected phyla. We combined data for five ecdysozoan phyla (Priapulida, Nematoda, Onychophora, Tardigrada and Arthropoda) and identified DNMT1, DNMT2 and DNMT3 in four out of these phyla. The only exception are Tardigrada, where neither DNMT1 and DNMT3 was detected. This suggests the absence of DNA methylation in, at least the currently sequenced, tardigrade species. Our data shows that DNA methyltransferases evolved independently and differently in the studied phyla of Ecdysozoa.

We proposed an adapted method (`MethMod`) to predict the DNA methylation status in a given species based on coding sequence (CDS) data. It was optimized over a wide phylogenetic range and requires only a single decisive parameter (the distance between the component means of a Gaussian Mixture Modelling) to achieve high specificity. Naturally, the method is limited if changes in the methylome have not yet altered the underlying genome significantly or if methylation is only present in small amounts. `MethMod` is available as a stand-alone python script and can be easily applied to emerging model organisms since only coding sequence data is required.

The data presented here will help to guide future projects to experimentally study DNA methylation in non-model Ecdysozoa species. The proposed analysis should be a worthwhile addition to newly sequenced genomes. It allows to expand their scope from the genomic to the epigenomic level.

## Declarations

### Funding

This work was supported by DFG STA 16/1 and 16/2 to PFS. JE was supported by Joachim Herz Stiftung.

### Conflicts

The authors have no conflict of interests to declare.

### Data Availability

The data underlying this article are available in the article and in its online supplementary material.

### Code Availability

`MethMod` which was used to predict the DNA methylation status is available on github: https://github.com/JanLeipzig/MethMod/

### Ethical approval

Not applicable.

### Consent to participate

Not applicable.

### Consent for publication

Not applicable.

## Notes

### Competing Interest Statement

The authors have declared no competing interest.

